# Cas Adaptor Proteins Coordinate Sensory Axon Fasciculation

**DOI:** 10.1101/159194

**Authors:** Tyler A. Vahedi-Hunter, Martin M. Riccomagno

## Abstract

Development of complex neural circuits like the peripheral somatosensory system requires intricate mechanisms to ensure axons make proper connections. While much is known about ligand-receptor pairs required for dorsal root ganglion (DRG) axon guidance, very little is known about the cytoplasmic effectors that mediate cellular responses triggered by these guidance cues. Here we show that members of the Cas family of cytoplasmic signaling adaptors are highly phosphorylated in central projections of the DRG as they enter the spinal cord. Furthermore, we provide genetic evidence that Cas proteins act DRG-autonomously to promote fasciculation of sensory projections. These data establish an evolutionarily conserved requirement for Cas adaptor proteins during peripheral nervous system axon pathfinding. They also provide insight into the interplay between axonal fasciculation and adhesion to the substrate.

## Introduction

Precise assembly of the peripheral somatosensory system involves migration of neural crest cells (NCCs) to coalesce into sensory ganglia and subsequent guidance of axonal projections from these newly formed ganglia. The NCCs that give rise to the dorsal root ganglia (DRG) originate from the dorsal spinal cord and migrate ventro-medially between the neural tube and rostral somite [1, 2]. Upon reaching the DRG, these neural progenitors continue to proliferate before committing to a neuronal or glial fate [3, 4]. The newly born sensory neurons then extend a central and a peripheral axon branch, acquiring the characteristic pseudounipolar morphology [5]. The resulting central projections traverse towards the spinal cord and enter the central nervous system (CNS) through the Dorsal Root Entry Zone (DREZ), while the distal processes navigate long distances to innervate their peripheral targets [6, 7]. Accurate guidance and fasciculation of these axons requires an intricately choreographed array of signaling cues acting on their cognate receptors [8, 9]. Although much is known about the ligand-receptor pairs required for axon trajectories, very little is known about the cytoplasmic effectors that allow these axons to respond to guidance cues.

Cas signaling adaptor proteins mediate a variety of biological processes including cell migration and changes in cell morphology [10], and exhibit specific expression patterns during neural development in rodents [11]. Cas proteins interact with various classes of signaling proteins, including cytosolic tyrosine kinases (like Src and Fak). Upon phosphorylation, Cas proteins can provide docking sites for SH2-containing effectors, including Crk, which stimulate Rac1-mediated actin remodeling [12]. We have recently uncovered an essential role for Cas family members during retinal ganglion cell migration [13], yet our current understanding of Cas adaptor protein function during mammalian axon pathfinding *in vivo* is limited. One member of this family, p130Cas, has been proposed as a required downstream component of netrin-mediated commissural axon guidance in the chicken spinal cord [14]. Interestingly, Drosophila Cas (dCas) has been shown to participate downstream of integrin receptors in axon fasciculation and guidance of peripheral motor axons [15]. Whether Cas proteins play similar roles during mammalian peripheral nervous system (PNS) development is currently unknown.

Here we examine the requirement for Cas adaptor proteins during DRG axon pathfinding. Our genetic data supports a novel role for Cas adaptor proteins during the fasciculation and guidance of central DRG projections in the DREZ. These data provide insight into the interplay between adhesion to the substrate and axon fasciculation.

## Results

Cas adaptor proteins have been shown to participate in the formation of the neuronal scaffold of the mammalian retina [13]. In addition, dCas is required for integrin-mediated peripheral axon guidance and fasciculation in Drosophila [15]. To investigate the role for Cas signaling adaptor proteins during mammalian PNS axon guidance, we started by assessing the expression pattern of *Cas* genes during embryonic development by *in situ* hybridization (Figure 1A-L). *Cas* genes are broadly expressed in the DRG and dorsal spinal cord at embryonic day (e)10.5 (Figure 1A-F). *p130Cas* and *Sin* continue to be expressed in the DRG at e12.5 and e14.5 (Figure 1 G, I, J, L). *CasL* DRG expression becomes weaker as development progresses, and becomes undetectable by e14.5 (Figure 1 B, E, H, K). All three *Cas* family members maintain expression in the dorsal spinal cord until at least e14.5 (Figure 1D-L). Sense negative control probes displayed negligible staining (Figure 1M-O; data not shown).

**Figure 1.**
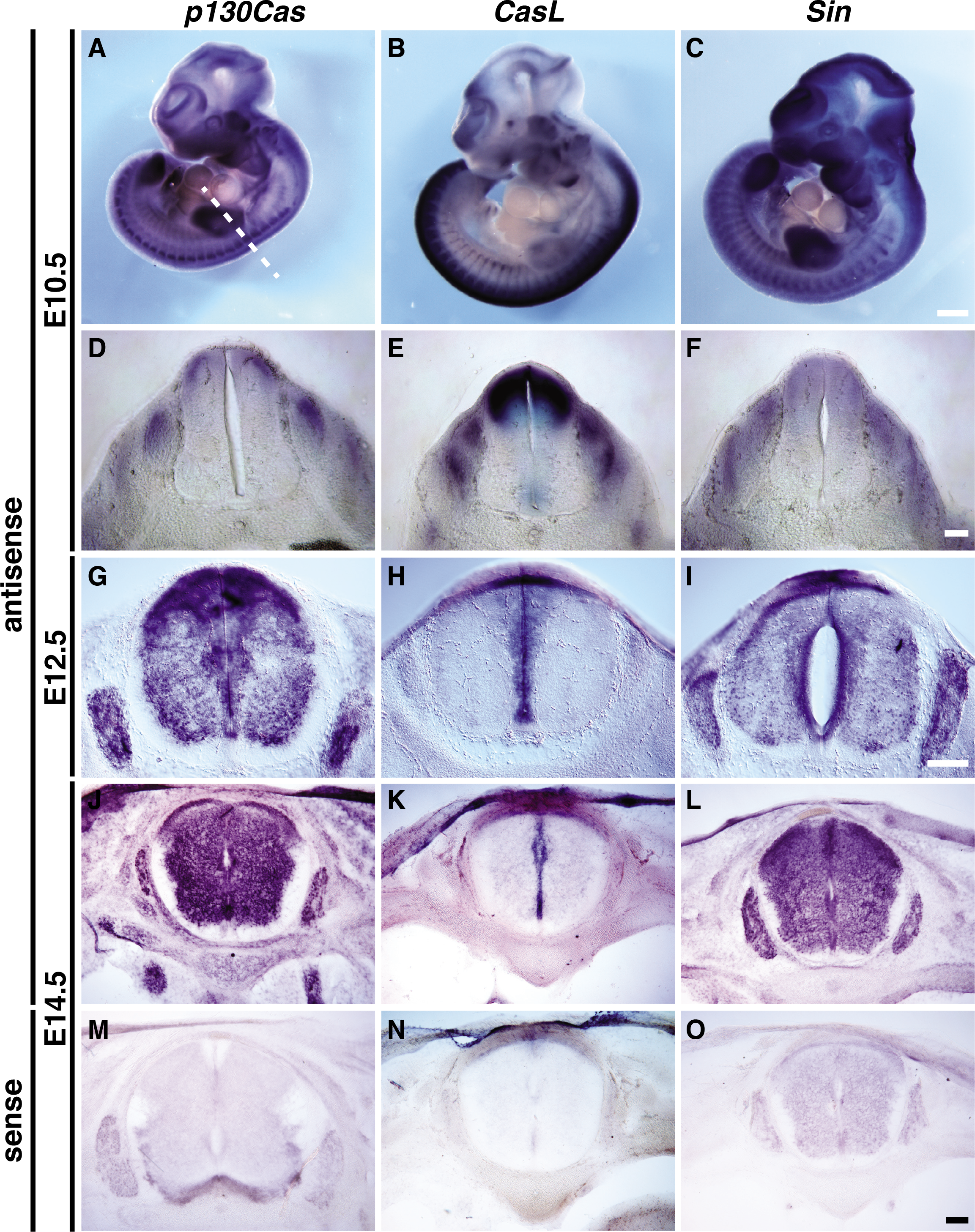
Expression of Cas *genes* in the developing dorsal root ganglia and spinal cord. (A-C) Whole-mount *in situ* hybridization for *p130Cas, CasL* and *Sin* in e10.5 embryos. (D-F) Transverse sections of embryos shown in A-C. Dotted line shows the plane of sectioning. (G-O) Transverse sections through embryonic spinal cords stained by in situ hybridization with probes against *p130Cas* (G, J, M), *CasL* (H, K, N) and *Sin* (I, L, O), at e12.5 (G-I) and e14.5 (J-O). *Cas* genes show overlapping expression in the DRGs and dorsal spinal cord. No staining was detected for the sense probes (M-O). Scale bars: 500 μm for A-C; 100 μm for D-F; 200 μm G-I and 100 pm J-O.

We next performed histological analyses of p130Cas protein expression in the developing spinal cord and DRG (Figure 2A). The expression pattern of p130Cas protein overlaps with that of *p130Cas* mRNA in DRG and spinal cord cell bodies (Figure 1G, 2A). In addition, p130Cas protein localizes to DRG central projections and spinal cord commissural axons, and is highly enriched in the DREZ and dorsal funiculus (Figure 2A). This expression was confirmed by utilizing a GENSAT BAC transgenic mouse line that expresses enhanced GFP (EGFP) under the control of *p130Cas* regulatory sequences [16, 17]. This transgenic line allows for the detection of cells expressing *p130Cas* [13]. The *p130Cas-EGFP-Bac* spinal cord EGFP expression pattern is consistent with that of endogenous p130Cas in wildtype (WT) animals (Figure 2A-C). As phosphorylation of Cas adaptor proteins mediates adhesion signaling during neural development [13, 15, 18], we examined the localization of Phosphotyrosine-p130Cas (PY-Cas) in the developing spinal cord and DRG. PY-Cas is present in the ventral funiculus and commissural axons as they reach the midline (Figure 2D, E). Interestingly, PY-Cas is also enriched in the DREZ, DRG central projections and dorsal funiculus (Figure 2D, F). Therefore, *Cas* mRNA, p130Cas protein, and phospho-tyrosine-p130Cas expression patterns are consistent with Cas involvement in DRG and commissural axon guidance.

**Figure 2.**
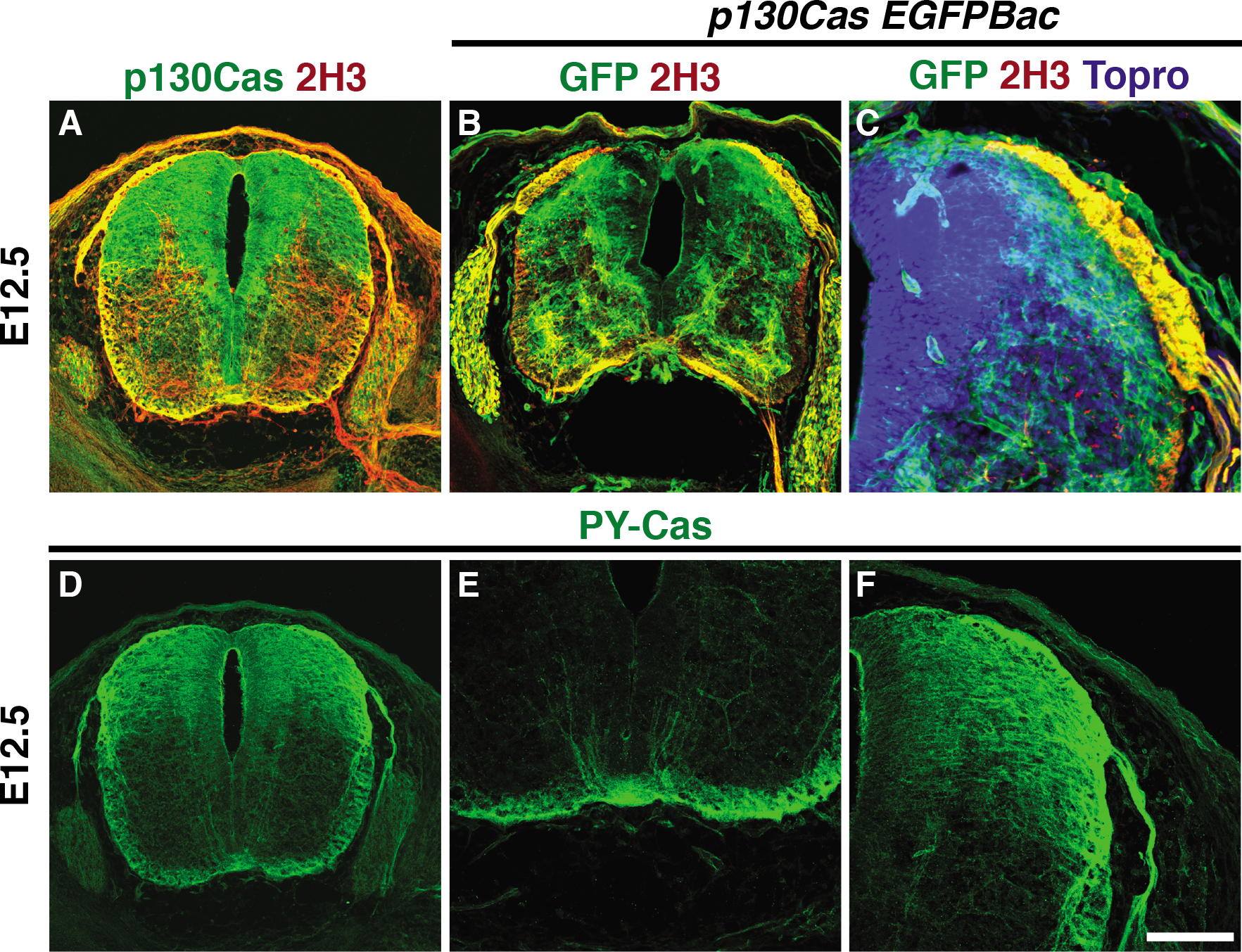
p130Cas is phosphorylated in commissural axons and DRG central projections. (A) Expression profile of p130Cas protein (green) in transverse sections through the mouse spinal cord at e12.5. Anti-Neurofilament (2H3, red) was used to reveal axons. (B-C) Immunofluorescence for EGFP (green) and 2H3 (red) in e12.5 *p130Cas EGFP-Bac* spinal cords. Topro (blue) was used to counterstain nuclei. (D-F) Expression of a phosophorylated-p130Cas (PY-Cas, green) in e12.5 spinal cord and DRGs. (E) PY-Cas is present in the ventral funiculus and commissural axons. (F) p130Cas phosphorylation is mainly found in the dorsal spinal cord and is enriched in DRG axons and DREZ. Scale bar: 200 pm for A, B, D and 100 pm for C, E, F.

Given the expression and phosphorylation pattern of Cas adaptor proteins in the developing spinal cord and DRG, we next asked whether *Cas* genes are required for DRG and commissural axon pathfinding. Since the expression patterns of *Cas* genes during spinal cord development are highly overlapping and Cas adaptor proteins act redundantly during retina development [13], we concurrently ablated all *Cas* genes from the dorsal spinal and DRG (dSC+DRG). Using *Wnt1Cre-2*a mice that expresses Cre recombinase in the dorsal spinal cord and neural-crest derived structures [19], we ablated a conditional allele of *p130Cas* in a *CasL*^-/-^;*Sin*^-/-^ double null mutant genetic background (we refer to *p130Cas^f/Δ^;CasL^-/-^;Sin^-/-^* mice as triple conditional knock-outs: “TcKO”) [13]. In control embryos, axons from DRG sensory neurons bifurcate and project along the anterior-posterior axis of the dorsal spinal cord as a tightly fasciculated bundle, as revealed by whole-mount immunohistochemistry at e12.5 (Figure 3A, B). This is in stark contrast to the DRG central projections of *Wnt1Cre;TcKO* embryos, which are highly defasciculated (Figure 3C-E). All other combinations of *Cas* family alleles display no overt phenotypes in DRG or other PNS axon tract guidance (data not shown). Interestingly, some of the defasciculated axons in *Wnt1Cre;TcKO* embryos appear to project towards the ventricular zone (Figure 3B, D-D’). These phenotypes observed in *Wnt1Cre;TcKO* embryos are 100% penetrant (n=6).

**Figure 3.**
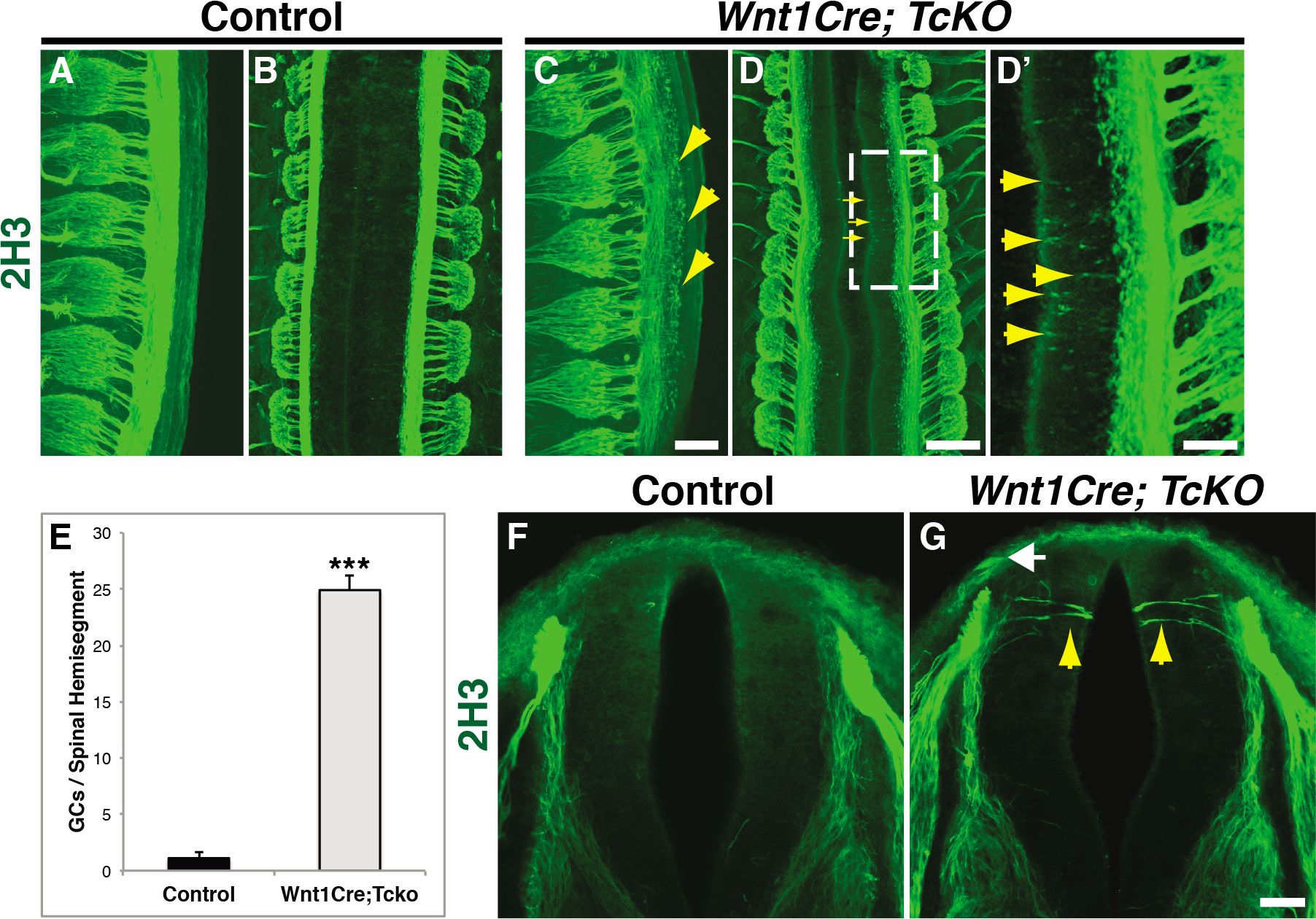
Cas adaptor proteins are required for the fasciculation of DRG central projections. (A-D’) Whole embryo immunostaining for neurofilament (2H3, green) at e12.5, from a side view (A, C) or a dorsal view (B, D, D’). D’ shows a higher magnification view of the doted area in D. The centrally projecting DRG axons are severely defasciculated as they enter the spinal cord. n=6 per gentoype; presented phenotypes displayed 100% penetrance. (**E**) Quantification of free growth cones (GCs) per spinal hemisegment, visualized from the side. (**F-G**) Transverse vibratome sections through e11.5 Control (F) and *Wnt1Cre; TcKO* (G) spinal cords at forelimb level stained using an antibody against neurofilament (2H3). Sensory axons invade the spinal cord gray matter prematurely in *Wnt1Cre; TcKO* animals (G). n=5 for each genotype; 100% penetrance. Scale bars: 100 μm for A, C; 200 μm B, D; 66.7 μm for D’; and 50 μm for F, G.

To further explore the role of Cas proteins during DRG axon pathfinding, we examined in more detail the innervation of the SC grey matter by sensory axons in control and *Wnt1Cre;TcKO* mutants. DRG afferent axons project to the DREZ and then stall during a “waiting period” before innervating the spinal cord. In the mouse this period extends from e11 until e13.5 for proprioceptors, or e15 for nociceptors [20, 21]. In control e11.5 embryos, no sensory axons are detected medial to the DREZ and dorsal funiculus (Figure 3F). However, several DRG axons invade the grey matter of *Wnt1Cre;TcKO* embryos prematurely (Figure 3G). This suggests that Cas adaptor proteins are required for proper fasciculation of DRG axons at the dorsal funiculus, as well as preventing these axons from entering the SC grey matter prematurely. Since Cas proteins are required for DRG axon fasciculation and guidance, we hypothesized that Cas proteins may be required for the guidance of other peripheral nerves. Could cranial nerves also require *Cas* gene function for proper fasciculation and guidance? Normally, vagal nerve central projections join and fasciculate with descending axonal tracks coming from the midbrain (Figure 4A) [22]. Interestingly, in *Wnt1Cre;TcKO*, the vagal afferents overshoot the descending midbrain tracks and display a defasciculated phenotype (Figure 4B). In addition to the vagal nerve, the trigeminal nerve also shows a distinct phenotype in *Wnt1Cre;TcKO* animals (Figure 4 C-F). Whereas the ophthalmic branch appears to be under-branched, the maxillary branch is highly defasciculated in *Wnt1Cre;TcKO* (Figure 4D, F) compared to control littermates (Figure 4C, E). Overall, these data support a role for Cas adaptor proteins during peripheral nerve pathfinding.

**Figure 4.**
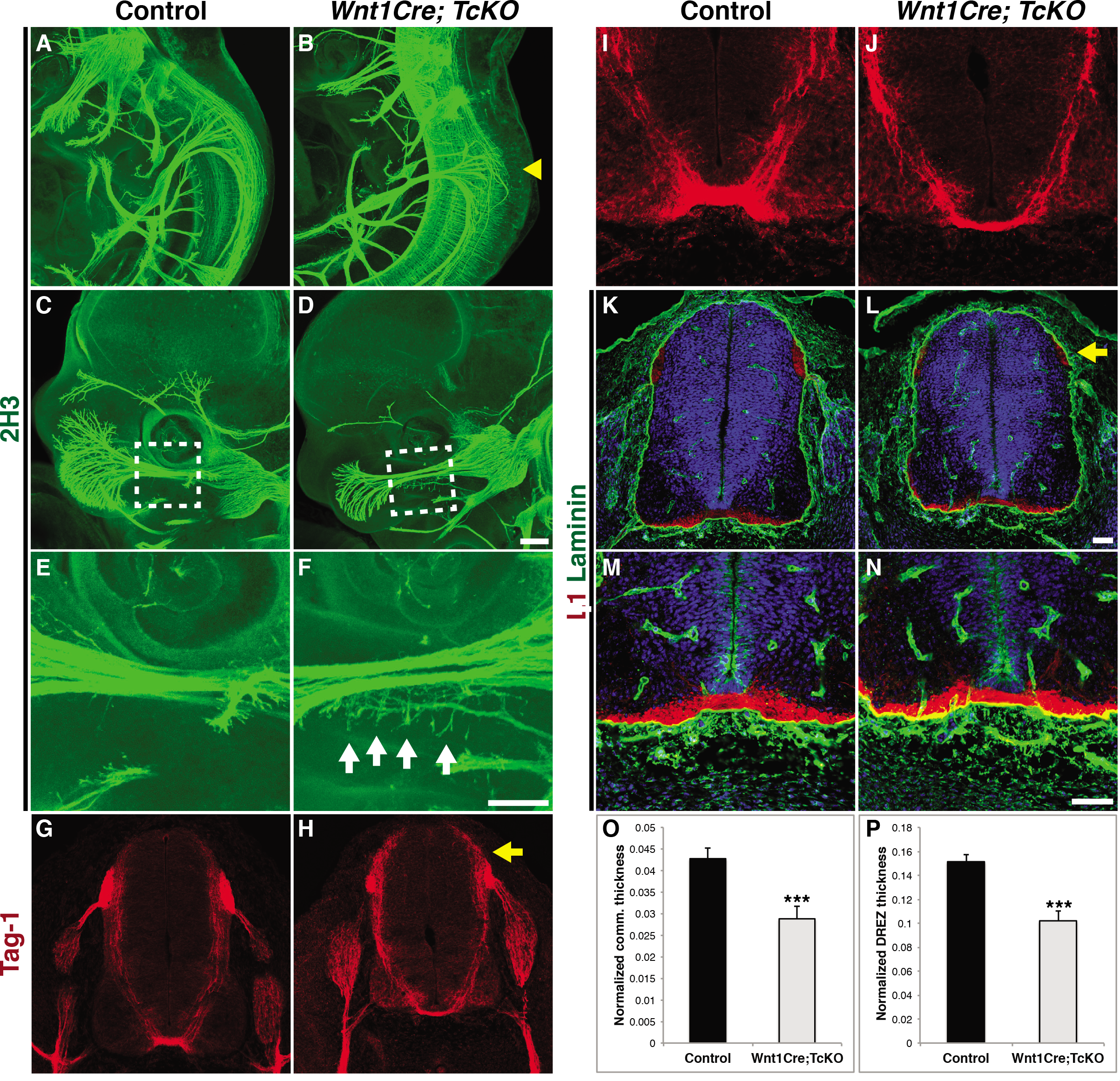
Cas adaptor proteins are essential for cranial nerve development, but mostly dispensable for commissural axon guidance. (A-H) Whole-mount immunostaining of Control (A, C, E) and *Wnt1Cre; TcKO* embryos (B, D, F) at e11.5 (A,B) and e12.5 (C-F). The central projections of the vagal nerve are severely defasciculated in *Wnt1Cre; TcKO* embryos (B, yellow arrowhead). The ophthalmic branch trigeminal nerve appears underbranched in *Wnt1Cre; TcKO*(D) embryos compared to controls (C). (E-F) Higher magnification view of white boxes in C and D reveals exuberant defasciculation of the maxillary branch of the trigeminal nerve in *Wnt1Cre; TcKO* embryos (F). n=6, 100% penetrance. (G-N) Transverse cryosections through control (G, I, K, M) and *Wnt1Cre;TcKO* (H, J, L, N) e11.5 spinal cords immunostained for Tag1 (G-J), L1Cam (K-N, red) and laminin (K-N, green). Sections in K-N were counterstained with Topro (blue). I-J, and M-N are higher magnification views of G-H, and K-L, respectively. The there is a reduction in the width of the ventral commissure (J, N), and the DREZ is smaller and disorganized in *Wnt1Cre; TcKO* embryos (yellow arrow, H, J). (O) Quantification of the normalized commissure thicknes in control and *Wnt1Cre;TcKO* embryos. (P) Quantification of the normalized DREZ thickness in control and *Wnt1Cre;TcKO* embryos. Scale bars: 200 μm for A-D; 100 μm for E, F; 50 μm for G,H K, L; and 50 for I, J, M, N

A previous report using small interference RNA (siRNA) knock-down suggested that *p130Cas* is required for commissural axon guidance [14]. We revisited this finding by taking advantage of our complete *Cas* loss of function mouse model (Figure 4G-O). We labeled commissural axons using the precrossing commissural axon marker Tag1 and the post-crossing marker L1 [23]. A significant reduction in the thickness of the ventral commissure was observed in *Wnt1Cre;TcKO* compared to control (Figure 4 G-O). This suggests that Cas genes might indeed play a conserved role in commissural axon guidance. Tag1 and L1 immunostaining also revealed a significant disorganization and reduction of the size of the DREZ in *Wnt1Cre;TcKO* animals compared to control (Figure 4G, H, K, L, P). These results illustrate the essential and multifaceted role of Cas proteins during both CNS and PNS circuit assembly.

Basement membrane (BM) integrity is required for proper axon guidance [22]. Thus, the abnormal fasciculation and guidance phenotypes observed at the DREZ in *Wnt1Cre;TcKO* embryos could be a secondary consequence of a disrupted basement membrane surface surrounding the spinal cord. To determine whether *Cas* genes are required for formation of the BM of the spinal cord, we analyzed its integrity in *Wnt1Cre;TcKO* animals. We visualized the BM using an antibody against laminin (Figure 4K-N). The BM appears to be intact in *Wnt1Cre;TcKO* embryos, and is indistinguishable from control embryos (Figure 4K-N). This suggests that *Cas* genes are dispensable for spinal cord BM formation, and that disruption of the basement membrane is unlikely to be responsible for axon pathfinding defects observed in *Wnt1Cre;TcKO* DRG central projections.

Selective ablation of *Cas* genes from the dorsal spinal cord and neural-crest derived PNS ganglia results in axon fasciculation and guidance defects (Figure 3 and 4); is there a DRG-autonomous requirement for Cas genes for axon pathfinding? To answer this question we took advantage of a transgenic line that expresses Cre recombinase under control of the human tissue plasminogen activator promoter (Ht-PA). This *HtPACre* transgene is expressed in migratory neural crest cells and their derivatives, including DRG, trigeminal and vagal ganglia [24], but not in the dorsal neural tube. We examined the DRG central and peripheral projections of control and *HtPACre; TcKO* embryos by whole-mount neurofilament immunofluorescence (Figure 5A-D). Interestingly, *HtPACre; TcKO* animals (Figure 5B) display aberrant defasciculation of DRG central projections compared to control littermates (Figure 5A, E). This abnormal defasciculation phenotype looks strikingly similar to that of *Wnt1Cre;TcKO* embryos (Figure 3A-E), suggesting that Cas genes act DRG-autonomously during central projection fasciculation and guidance. However, *HtPACre*; *TcKO* vagal and trigeminal nerve projections look indistinguishable from controls (data not shown), raising the possibility that Cas genes might act in a non-autonomous manner during the fasciculation of these two nerves.

**Figure 5.**
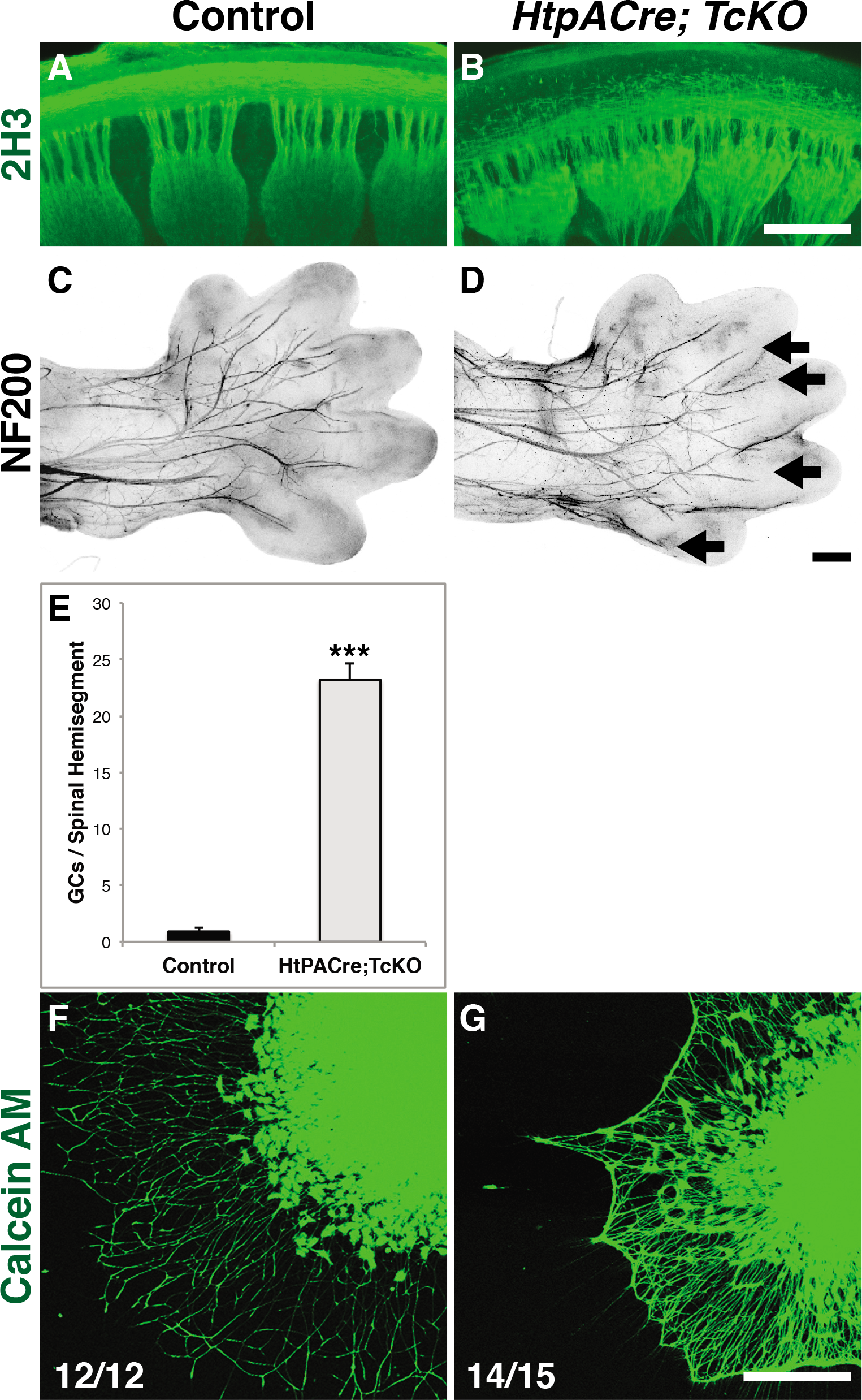
DRG-autonomous requirement for *Cas* genes. (A-D) Wholemount neurofilament immunostaing of control (A, C) and *HtpACre; TcKO* embryos (B, D). Sideview of e12.5 spinal cords stained with 2H3. The defasciculation of DRG central projection observed in *HtpACre; TcKO* embryos (B) resembles that of *WntlCre; TcKO* embryos (Figure 3). N=4 for each genotype, 100% penetrance. (C, D) Dorsal view of e14.5 limbs stained with NF200. Select axonal branches that innervate the digits appear hyperfasciculated in *HtpACre; TcKO* (D) hindlimbs (white arrows). Fluorescent images were inverted to facilitate visualization. n=6 limbs per genotype. (E) Quantification of free growth cones (GCs) per spinal hemisegment, visualized from the side. (F-G) DRG explants from control (F) and *HtpACre; TcKO* (G) embryos. Cells were visualized using Calcein-AM. *HtpACre; TcKO* explants display an abnormal “cobweb” morphology (G). Scale bars: 200 μm for A, B; 250 μm for C, D and 200 μm for F, G.

The stereotyped innervation pattern of the limb by sensory axons provides an excellent model to analyze DRG peripheral projection branching and fasciculation [25, 26]. The early lethality of *Wnt1Cre;TcKO* animals (between e12 and e13) precluded us from performing analysis of limb innervation in those animals. Because *HtPACre*; *TcKO* animals survive at least until early adulthood, we explored hind-limb innervation in *HtPACre*; *TcKO* e14.5 embryos and control littermates using neurofilament 200 (NF200), a marker for mechanosensory aβ fibers (Figure 5C, D). The innervation pattern in *HtPACre*; *TcKO* hind-limbs is abnormal compared to control limbs (Figure 5C, D). Mechanosensory fibers stall prematurely and hyper-fasciculate in *HtPACre*; *TcKO* animals. These results demonstrate a requirement of *Cas* genes in a DRG-autonomous manner for the guidance and fasciculation of somatosensory peripheral and central projections.

Based on our results, we hypothesized that Cas adaptor proteins regulate fasciculation at choice-points by allowing the growing axons to distinguish between adhesion to the extracellular matrix (ECM) and to other axons. To investigate how Cas adaptor proteins participate in the fasciculation and defasciculation of axons at choice-points, we cultured e13.5 DRG explants from control and *HtPACre*; *TcKO* on the ECM protein laminin. This provides a very simple model to test for the ECM vs. axon adhesion choice. Control DRG explants axons grew radially and display a characteristic sun-like morphology (Figure 4F). Interestingly, although the initial axonal growth of *HtPACre; TcKO* appears normal, the axons fasciculate together forming a rim of axon bundles at a distance from the explant (Figure 4G). This “cobweb” phenotype was never observed in control DRG explants under the same conditions (Figure 4F). These data indicate that Cas proteins are necessary for balancing DRG axon fasciculation and defasciculation *in vivo* and *in vitro.*

## Discussion

Here, we demonstrate an evolutionarily conserved requirement for Cas adaptor proteins during guidance and fasciculation of PNS axons. Our results in the mouse are consistent with the previously described role of *dCas* in Drosophila PNS development [15]. In Drosophila, Cas phosphorylation and function during PNS axon fasciculation and guidance is mainly regulated by integrins. Similarly, during mammalian retina migration, integrin-β1 appears to be the primary regulator of Cas function, as shown by the identical phenotype in their respective null mutants, and molecular epistasis results [13]. Whereas the peripheral innervation phenotypes in the developing limb are similar in *HtPACre;Itgb1*^*f/f*^ and *HtPACre;TcKO* mutants, the severe central projection phenotype observed in *Cas TcKO* mice was not observed in their *mtegrin-β1* counterparts [25]. This suggests that integrin-β1 is not the sole upstream regulator of Cas adaptor function during DRG central projection pathfinding. Whether integrins act redundantly to regulate Cas function or a different guidance cue receptor is involved in this process remains to be investigated. Interestingly, *HtPACre*; *TcKO* vagal nerve central projections look indistinguishable from controls, suggesting that Cas genes might act in a non-autonomous manner during the fasciculation of these projections. It is also possible that, because vagal projections fasciculate with descending midbrain projection, expression of *Cas* just in the midbrain projections is enough to guarantee fasciculation of the two tracks.

In regards to Cas function during commissural axon guidance, it was reported that *p130Cas* mediates Netrin signaling during this pathfinding event [14]. The single *Wnt1Cre;p130Cas^f/f^* mutants displayed no obvert axon guidance phenotypes (data not shown). The complete *Cas* loss-of-function mouse model *(Wnt1Cre;TcKO)* did show a significant thinning of the commissure by e11.5, although the phenotypes observed in the chicken knock-down experiments were much more striking [14]. Furthermore, the observed phenotype was notably milder than that of the *Netrin^-/-^* mice, which have almost no detectable ventral commissures [27]. This result suggests that if *Cas* genes indeed act downstream of Netrin during mouse commissural axon guidance, they would more likely serve as modulators than obligate-downstream effectors. Another possibility is that *Cas* genes might play a more general role during commissural axon fasciculation.

An unexpected discovery was the fact that *Cas-null* DRG axons displayed a different growth pattern on laminin than control explants, re-fasciculating with each other at a distance from the somas (Figure 5F, G). This result raises the intriguing possibility that *Cas* genes are required for neurons to distinguish between secreted adhesion molecules in the ECM (e.g. laminin) and neural adhesion molecules present in axons themselves. This environmental assessment will be particularly important at choice-points like the DREZ, and could offer a potential mechanism underlying the DRG central projection defasciculation phenotypes observed in *HtpACre*; *TcKO* and *Wnt1Cre;TcKO* embryos. Alternatively, Cas might be important for DRG axons to pause at the DREZ to sense repulsive and attractive cues on their way to finding their targets. In this regard it is interesting to note that some of the sensory phenotypes observed in *HtpACre*; *TcKO* and *Wnt1Cre;TcKO* embryos resemble aspects of *Robo/Slit* [28, 29], *dystroglycan* [22], netrin [30], and *Neuropilin-1* [20] mutants. Future studies will investigate the role of Cas adaptor proteins during the interplay between adhesion to the substrate and guidance cue signaling.

## Materials and methods

### Animals

The day of vaginal plug observation was designated as embryonic day 0.5 (e0.5) and the day of birth postnatal day 0 (P0). Generation of the *HtpACre, Wnt1Cre, p130CaS*^*f/f*^, *CasL^-/-^* and *Sin^-/-^* transgenic mouse lines has been described previously [13, 19, 24, 31, 32].

### *In situ* Hybridization

*In situ* hybridization was performed on spinal cord frozen sections (20 μm thickness) using digoxigenin-labeled cRNA probes, as previously described [33]. Whole-mount RNA in situ hybridization was performed as described [34]. Generation of the *p130Cas, CasL* and *Sin* cRNA probes has been described in [13].

### Immunofluorescence

Mice were perfused and fixed with 4% paraformaldehyde for 1 hour to O/N at 4°C, rinsed, and processed for whole-embryo staining or sectioned on a vibratome (75 *μ*m). Whole-mount immunofluorescence was performed as described in [35]. Immunohistochemistry of floating sections was carried out essentially as described [36]. For cryostat sections, following fixation, embryos were equilibrated in 30% sucrose/PBS and embedded in OCT embedding media (Tissue-Tek). Transverse spinal cord sections (20-40 *μ*m) were obtained on a Leica CM3050 cryostat and blocked in 10% goat serum in 1 X PBS and 0.1% Triton-X100 for 1 hr at room temperature. Sections were then incubated O/N at 4°C with primary antibodies.

Sections were then washed in 1 X PBS and incubated with secondary antibodies and TOPRO-3 (Molecular Probe at 1:500 and 1:2000, respectively). Sections were washed in PBS and mounted using vectorshield hard-set fluorescence mounting medium (Vector laboratories). Confocal fluorescence images were taken using a Leica SPE II microscope. Primary antibodies used in this study include: rabbit anti-p130Cas C terminal (Santa Cruz, 1:200), rabbit anti-p130Cas PY165 (Cell Signaling Technology, 1:100), rabbit anti-laminin (Sigma, 1:1000), rabbit anti-GFP (Lifescience Technologies, 1:500), chicken anti-GFP (AVES, 1:1000), mouse anti-Neurofilament (2h3, Developmental Studies Hybridoma Bank, 1:500), mouse anti-Tag1 (4D7, Developmental Studies Hybridoma Bank, 1:50), Rat anti-L1 (MAB5272, Millipore, 1:500) and rabbit anti-NF-200 (Millipore, 1:500).

### Quantification of spinal cord ventral commissure and DREZ thickness

Thickness of the DREZ and ventral commissure were measured on L1-immunostained cryosections at e11.5 (20-μm sections). The thickness values for the ventral commissure were normalized to the distance between roof plate and floor plate for each section, as described previously [37, 38]. The maximal thickness of the DREZ for each hemi-spinal cord was recorded and normalized to the distance between the BM and the ventricular zone at the same dorso-ventral level. Thickness was measured at branchial levels. Five sections per embryo, from 3-5 embryos were analyzed. Statistical differences for mean values between two samples were determined by two-tail Student’s t-test for independent samples.

### Tissue Culture

DRGs from e13.5 embryos were dissected in ice-cold L15 (Invitrogen). DRG explants were then plated on acid-washed glass coverslips previously coated with 5μg/ml laminin and 100μg/ml polyD-lysine. DRGs were then cultured for 18 hours in enriched Opti-MEM/F12 media containing 15ng/ml NGF, as previously described [39]. Live explants were stained with Calcein-AM (Invitrogen) and then imaged. A total of 12-15 explants from 3 independent experiments were analyzed.

## List of Abbreviations

**NCC** - neural crest cell

**DRG** - dorsal root ganglia

**CNS** - central nervous system

**DREZ** - dorsal root entry zone

**Cas** – Crk-associated substrate

**Src** - sarcoma

**Fak** – focal adhesion kinase

**SH2** – sarcoma homology domain 2

**Crk** – cdc2-related protein kinase

**dCas** - Drosophila Cas

**PNS** - peripheral nervous system

***Sin*** - sarcoma-interacting Cas

***CasL*** - Cas in lymphocyte

**p130Cas** - 130 KDa Crk-associated substrate

**BAC** - bacterial artificial chromosome

**GFP** - green fluorescent protein

**WT** – wild-type

**PY-Cas** - phosphotyrosine-p130Cas

**dSC** - dorsal spinal cord

**TcKO** - triple conditional knock-out

***Wnt1*** – wingless-related integration site 1 (gene)

**SC** – spinal cord

**siRNA** - small interference RNA

**Tag1** – neuronal marker

**L1** – L1 cell adhesion molecule

**BM** - basement membrane

**(Ht-PA)** - human tissue plasminogen activator promoter

**NF200** - neurofilament 200

**ECM** - extracellular matrix

***Itgb1*** – integrin-β1(gene)

**cRNA** – copy ribonucleic acid

**PBS** – phosphate buffer saline

**OCT** – optimum cutting temperature

**NGF** – neuronal growth factor

## Declarations

### Acknowledgements

We thank Drs. Christopher Deppmann, Garret Anderson and Randal Hand for helpful comments on the manuscript.

## Availability of data and materials

All data analyzed during this study are included in this article.

## Authors Contributions

TAVH performed experiments and wrote the manuscript. MMR conceived the project, designed and performed experiments, and wrote the manuscript.

## Consent for publication

Not applicable.

## Ethics approval

All animal procedures presented here were performed according to the University of California, Riverside’s Institutional Animal Care and Use Committee (IACUC)-approved guidelines.

